# Proximity Labeling Expansion Microscopy (PL-ExM) resolves structure of the interactome

**DOI:** 10.1101/2023.11.09.566477

**Authors:** Sohyeon Park, Xiaorong Wang, Xiangpeng Li, Xiao Huang, Katie C. Fong, Clinton Yu, Arthur A. Tran, Lorenzo Scipioni, Zhipeng Dai, Lan Huang, Xiaoyu Shi

## Abstract

Elucidating the spatial relationships within the protein interactome is pivotal to understanding the organization and regulation of protein-protein interactions. However, capturing the 3D architecture of the interactome presents a dual challenge: precise interactome labeling and super-resolution imaging. To bridge this gap, we present the Proximity Labeling Expansion Microscopy (PL-ExM). This innovation combines proximity labeling (PL) to spatially biotinylate interacting proteins with expansion microscopy (ExM) to increase imaging resolution by physically enlarging cells. PL-ExM unveils intricate details of the 3D interactome’s spatial layout in cells using standard microscopes, including confocal and Airyscan. Multiplexing PL-ExM imaging was achieved by pairing the PL with immunofluorescence staining. These multicolor images directly visualize how interactome structures position specific proteins in the protein-protein interaction network. Furthermore, PL-ExM stands out as an assessment method to gauge the labeling radius and efficiency of different PL techniques. The accuracy of PL-ExM is validated by our proteomic results from PL mass spectrometry. Thus, PL-ExM is an accessible solution for 3D mapping of the interactome structure and an accurate tool to access PL quality.

## INTRODUCTION

Most cellular functions are realized by a set of protein-protein interactions (PPIs) called the protein interactome. Studies on the interactome of a hub protein transform our understanding of health and diseases and aid in discovering therapeutic targets^1-4^. Recent advancements in microscopy significantly advanced our understanding of protein interactomes by providing structural information from atomic to organellar scales. Cryo-electron microscopy uncovers atomic details of interacting proteins that predict binding sites. Super-resolution microscopy reveals molecular details that provide spatial relationships between specific interacting proteins. Scanning electron microscopy maps the overall proteome distribution which provides a global landscape of PPIs. Yet, visualization of the 3D architecture for the interactome has lagged^4,5^.

Visualizing the structural context of PPIs is essential for understanding how PPIs are organized by protein assembly and influenced by their subcellular environment. For example, by locating the activation of extracellular-signal-regulated kinase (ERK) by G-protein-coupled receptors (GPCRs) at endosomes, Kwon et al. identified a non-canonical mechanism of spatial regulation of ERK signaling through endosomal signaling^5^. Pownall et al. used ChromExM of embryos to reveal how the pioneer factor Nanog interacts with nucleosomes and RNA polymerase II (Pol II), providing direct visualization of transcriptional elongation as string-like nanostructures. The structural information of the interactome can enable us to discover new PPI mechanisms. There is an urgent need for imaging methods that can dissect the spatial relationships in the interactome.

Capturing the 3D architecture of the interactome presents a dual challenge: precise interactome labeling and super-resolution imaging. Precise interactome labeling should highlight the interactome of a targeted protein from the whole proteome of a cell. Proximity labeling (PL) emerged as a powerful technique that spatially selects proteins within its labeling resolution from the protein of interest. In this method, the protein of interest is fused to or labeled by an enzyme. When activated, this enzyme modifies nearby proteins by attaching a small marker like biotin to them. Proximity-labeled proteins can be subsequently analyzed by mass spectrometry (PL-MS) as potential interaction partners with the protein of interest. Several proximity labeling methods, such as HRP^6-8^, APEX^9-11^, BioID^12-14^, TurboID^15, 16^, and µMap^1^ have been widely used with mass spectrometry (MS) to identify the organellar proteome^10, 17^ and network of interactions in cells^14, 18-20^. These PL methods paved the way for interactome visualization by precisely labeling the interactome.

The second challenge in interactome visualization is simultaneously imaging specific proteins and its interactome structure with super resolution. Although super-resolution light microscopy can specify proteins and electron microscopy can visualize proximity-labeled proteins, it is difficult to simultaneously resolve both with the matching resolution. An emerging super-resolution technique called expansion microscopy (ExM) raised a promising solution. ExM is a chemical approach to increase the resolving power of any microscope by physically expanding cells by 4-20 times in each dimension ^21^. The early versions of ExM methods use antibodies and fluorescent proteins to label proteins, which only allow targeted protein imaging^22-24^. Excitingly, recent advances in ExM enabled super-resolution imaging of nonspecifically labeled biomolecules as the context channel in addition to the immunostained specific proteins. For instance, Mao et al. and M’saad et al. respectively demonstrated the power of their FLARE^25^ and pan-ExM^26^ methods in imaging the entire protein, lipid, and carbohydrate landscape. In another study, Pownall and colleagues mapped chromatin with single-nucleosome resolution using their technique chromExM^27^. Klimas et al. developed a Magnify protocol that retains nucleic acids, proteins and lipids in a uses a mechanically sturdy gel^28^. Beyond protein and DNA landscape, Sun et al. developed click-ExM enabling imaging of all biomolecules including glycans and small molecules^29^. These approaches collectively spotlight the ability to delineate specific proteins within context structures at a matching super-resolution. However, a glaring gap persists as ExM has not yet been used in studying the interactome, underscoring an unaddressed demand in interactome visualization.

We report proximity labeling expansion microscopy (PL-ExM), which simultaneously images the 3D architecture of the interactome and specific interactive proteins with super-resolution (Figure 1A). PL-ExM uses PL to label the interactome, antibodies to specify proteins of interest, and ExM for super-resolution imaging. The advantage of ExM over super-resolution light microscopy, such as STORM and STED, is its fast speed, high imaging depth, and low requirement for advanced microscopes. Using PL-ExM, we can locate specific proteins on the 3D structure of their interactome with a resolution up to 12 nm on commonplace microscopes, such as confocal and Airyscan. PL-ExM is compatible with any PL methods that can biotinylated proteins, for example, APEX and HRP labeling. Interestingly, HRP-catalyzed tyramide signal amplification (TSA) was recently used to amplify signals for ExM ^30^, but not for interactome visualization. PL-ExM was designed and optimized for the opposite purpose, that is proteome characterization.

**Figure 1.**
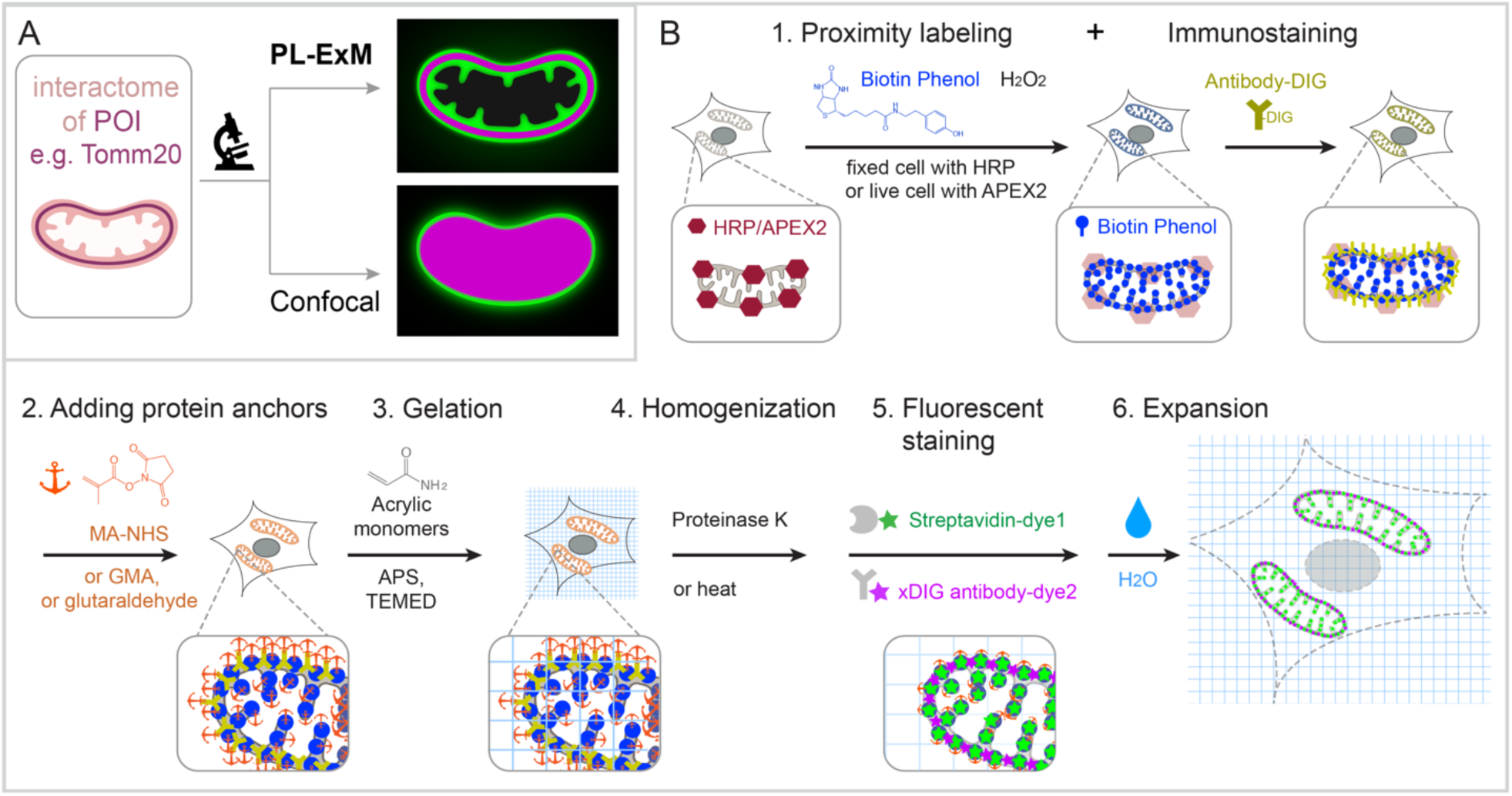
Graphic abstract and workflow of PL-ExM. In the showcase, Tomm20 is the bait for the PL and the target for the immunostaining. (A) Graphic abstract of PL-ExM method. PL-ExM offers super resolution to visualize small interactome structures that present the ground truth. Diffraction-limited microscopy, such as confocal microscopy, misses structural details in the ground truth. (B) The PL-ExM workflow comprises six steps. 1. Proximity labeling catalyzed by enzymes (HRP, APEX, etc.) and delivered by biotin phenol. Following PL, a protein of interest is labeled with antibodies conjugated with DIG. 2. Adding protein anchors, such as MA-NHS, GMA or glutaraldehyde. 3. Gelation with acrylic and acrylate monomers. 4. Denaturation using proteinase K or heat denaturation. 5. Fluorescent staining: stain the biotin and DIG with fluorescently conjugated streptavidin and anti-DIG antibodies. 6. Expansion: expand hydrogel through immersion in pure water.

Beyond imaging, this method can assess the quality of PL. Despite its importance of PL, the variability in the labeling resolution and efficiency of PL experiments often leads to limited overlap in PL-MS results, even when analyzing the interactome of the same targeted protein^31^. For example, a study showed less than 25% overlap in interactomes detected by APEX2 and BioID for the same bait valosin-containing protein (VCP) ^19^. Oakley et al. observed a 5-fold difference in labeling radius between µMap and peroxidase-based PL using STED^32^. Using PL-ExM, we compared the labeling radius and efficiency between APEX2 and HRP labeling, and between various labeling durations. To validate the PL-ExM imaging in evaluating PL quality, we profiled the interactome using PL-MS in parallel. The agreement between our imaging and MS data confirms that PL-ExM is a reliable and accurate tool for PL quality control.

We will unfold the workflow of PL-ExM and demonstrate its capability of interactome visualization and PL assessment as follows.

## RESULTS

### Principle and workflow

PL-ExM provides super-resolution to dissect the 3D architecture of the interactome by physically expanding the proximity-labeled cells and tissues in the swellable hydrogel. The effective imaging resolution of an expanded sample is equal to the microscope resolution divided by the length expansion factor of the sample. PL-ExM is compatible with most light microscopes, such as confocal, Airyscan, light sheet, SIM, STORM, and STED, and most ExM protocols which result in different expansion factors. For example, if the proximity labeled sample is expanded by four times and imaged with a confocal with a resolution of 280 nm, the effective imaging resolution will be 70 nm.

The swellable hydrogel that is made of different recipes and expansion procedures can expand from 3 to 14 times. The most commonly used gel formula for expansion microscopy consists of acrylamide, sodium acrylate, ammonium persulfate (APS), N,N,N′,N′-Tetramethylethylenediamine (TEMED), and N-N′-methylenebisacrylamide ^24, 33, 34^. This hydrogel expands about 4 times in pure water. By adjusting the crosslinkers or hydrolysis duration, the hydrogel can expand up to 13 times in one round ^35-39^. Multiple rounds of expansion even achieve a length expansion factor of 15 to ∼20x ^40^. The sample expansion improves the resolving power of the microscope by a factor from 3 to 20 depending on the expansion protocol. With different combinations of the microscope and the expansion protocol, PL-ExM achieves super resolution ranging from 12 nm to 70 nm, allowing visualization of a burst of structural details in the interactome that was not resolvable by diffraction-limited microscopes alone (Figure 1A).

The workflow of PL-ExM includes 6 steps (Figure 1B): 1. PL and immunostaining, 2. adding protein anchors, 3. gelation, 4. homogenization, 5. fluorescent staining, and 6. expansion. Technically, any PL method can be used as step 1. Peroxidase-based PL of mitochondria is showcased in our workflow because it is widely used. Peroxidase HRP or APEX2 is first introduced to bait protein of the interactome. In the presence of hydrogen peroxide (H2O2) and biotin-phenol, proteins within a labeling radius of the peroxidase are biotinylated. Additionally, a protein of interest is immunostained by antibodies conjugated with digoxigenin (antibody-DIG). Following the PL and immunostaining is the expansion procedure consisting of steps 2 to 6. In Step 2, proteins are chemically modified with anchoring molecules, such as glutaraldehyde (GA), methacrylic acid N-hydroxysuccinimide ester (MA-NHS), or glycidyl methacrylate (GMA). These anchors serve the same goal: covalently crosslinking proteins to polyacrylic chains when polyacrylic hydrogel is formed inside and outside of the cells in Step 3. Next, cells that are embedded in the hydrogel are homogenized by proteinase K digestion or heat denaturation (Step 4). The homogenization breaks the protein interactions to allow isotropic sample expansion in the final expansion step (Step 6). Before expansion, the biotinylated interactome and DIG-labeled proteins of interest are stained by fluorescently conjugated streptavidin and anti-DIG antibodies, respectively (Step 5). The reason to introduce fluorescent dyes after gelation is that free radical polymerization reactions can significantly quench fluorescent dyes ^23, 24, 34, 36, 41, 42^. We have demonstrated that post-gelation fluorescence staining of biotin or DIG probes can increase the signal-to-noise ratio of ExM images by several folds in our Label-Retention Expansion Microscopy (LR-ExM) technique^34^. Through the 6 steps, PL, ExM, and LR-ExM are streamlined into one workflow of PL-ExM.

Detailed chemical reactions underlining each step in the workflow are described in Figure S1.

### PL-ExM provides super resolution to visualize the 3D interactome architecture

We demonstrated the resolution improvement of PL-ExM by comparing the images of proximity-labeled mitochondria with and without expansion (Figure 2). The bait protein is the outer mitochondrial membrane (OMM) protein TOMM20, which was immunostained with antibodies conjugated with HRP. Proteins within the labeling radius of HRP were biotinylated by biotin-phenol in the presence of hydrogen peroxide. The PL duration was 30 seconds. The TOMM20 was also immunostained with antibody-DIG as the second color channel. Both expanded and non-expanded samples were imaged with the same Airyscan microscope, which has a measured resolution of 180 nm (Figure S2). Since this resolution was much larger than the labeling radius of HRP, images of non-expanded samples failed to encapsulate the intricate details of the mitochondria (Figure 2 A, B-E). On the contrary, PL-ExM imaging of 4.2 times expanded samples resolved the hollow structure of mitochondria (Figures 2J) and sometimes the mitochondria cristae (Figures S3B&S4B) with its 43 nm effective resolution. This observation of the hollow structure with high signal at the periphery and a medium signal inside (Figure 2J) indicated that the HRP proximity labeling of TOMM20 not only biotinylated proteins on the outer mitochondrial membrane, such as translocases of the outer membrane (TOMs), but also the ones inside, such as translocases of the inner membrane (TIMs). The protein identities are confirmed in our PL-MS analyses (Figure 4P).

**Figure 2.**
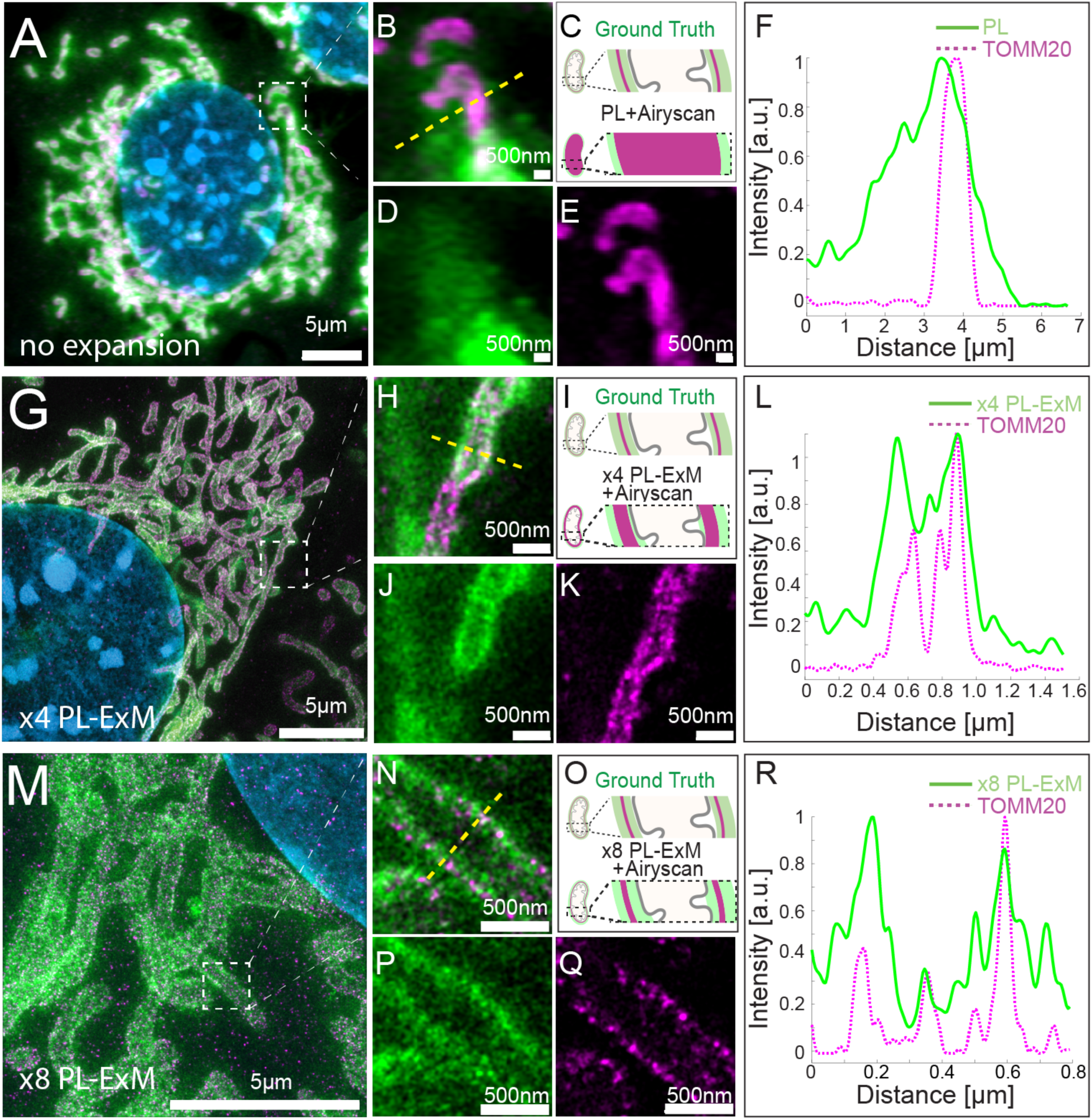
PL-ExM offers super resolution for the visualization of the proximity-labeled interactome landscape. All images were taken on MEF cells labeled with two colors in the same way. The TOMM20 was proximity-labeled to show its interactome (green) and simultaneously immunostained to locate the protein of interest (magenta). The nucleus was stained with DAPI (blue). All images were taken with Airyscan microscope. (A) Representative image of a non-expanded sample. (B) Magnified view of the boxed region in (A). (C) Schematics of the ground truth structure of proximity-labeled TOMM20 (green) and immunostained TOMM20 (magenta), and the expected image without expansion. (D) PL channel of (B). (E) Immunostained TOMM20 channel of (B). (F) A representative histogram showing the fluorescence intensity in a cross section of a mitochondrion from the image (B) of the non-expanded sample. The fluorescence intensity was denoised and normalized with respect to each channel. (G) Representative PL-ExM image of a 4-time expanded sample, named x4 PL-ExM. (H) Magnified view of the boxed region in (G). (I) Schematics of the same ground truth as in (C), and the expected PL-ExM image of the 4-time expanded sample. (J) PL-ExM channel of (H). (K) Immunostained TOMM20 channel of (H). (L) A representative histogram showing the fluorescence intensity in a cross section of mitochondrion from a x4 PL-ExM image. (M) Representative PL-ExM image of an 8-time expanded sample, named x8 PL-ExM. (N) Magnified view of the boxed region in (M). (O) Schematics of the same ground truth as in (C), and the expected PL-ExM image of the 8-time expanded sample. (P) PL-ExM channel of (N). (Q) Immunostained TOMM20 channel of (N). (R) A representative histogram showing the fluorescence intensity in a cross section of mitochondrion from an x8 PL-ExM image. In all histograms (F,L&R), the fluorescence intensity was denoised and normalized with respect to each channel. (A, G, M, N, P, Q) are maximum intensity projections of z stacks. (B, D, E, H, J, K) are single-slice images of 3D z stacks. Length expansion factors are 4.2 for samples (G, H, J, K), and 8.2 for (M, N, P, Q). All scale bars are in pre-expansion units.

The resolution of PL-ExM can be further improved with larger expansion factor. We expanded proximity-labeled cells by 8.2 times using the TREx protocol ^38^. As a result, x8 PL-ExM provided 22 nm resolution, which resolved two narrow and well-separated peaks of proximity-labeled proteins at the cross-section of mitochondrion (Figures 2P&R). The distance between the two peaks showed that the mitochondrion had a diameter of 500 nm (Figure 2R). The full width of the half maximum (FWHM) of each peak represented a PL resolution of 0.37µm (Figure 2R). In summary, PL-ExM can significantly increase the effective imaging resolution by 4 to 8 times with a single round of expansion.

### Multiplex Imaging reveals spatial relationships between interactive proteins

In the previous section, we used two-color PL-ExM to visualize the spatial relationship between the bait protein TOMM20 in its mitochondrial interactome. In this section, we demonstrated how to identify other interactive proteins in the interactome using the same method, with the following two examples.

Previous studies suggested that clathrin-coated pits (CCPs) are transported on microtubules based on live cell imaging ^43, 44^. Here, we try to confirm the CCP-microtubule interactions by directly locating CCPs in the microtubule interactome. We imaged immunostained Clathrin A (CLTA) and proximity-labeled α-TUBULIN using two-color PL-ExM (Figures 3A-G). Thanks to the super resolution, the images show that the proximity-labeled proteins not only displayed the microtubules but also showed clusters budding from the microtubules (pointed by arrows in Figures 3C&F). Interestingly, many of these clusters were found to be partially overlapping with the immunostained CCPs (Figures 3B&E). This is a direct visualization of CCPs as components of the interactome of microtubules, which affirms that CCPs interact with microtubules. Such spatial relationships in interactomes were not detectable without expansion due to limited resolution (Figures 3H-N).

**Figure 3.**
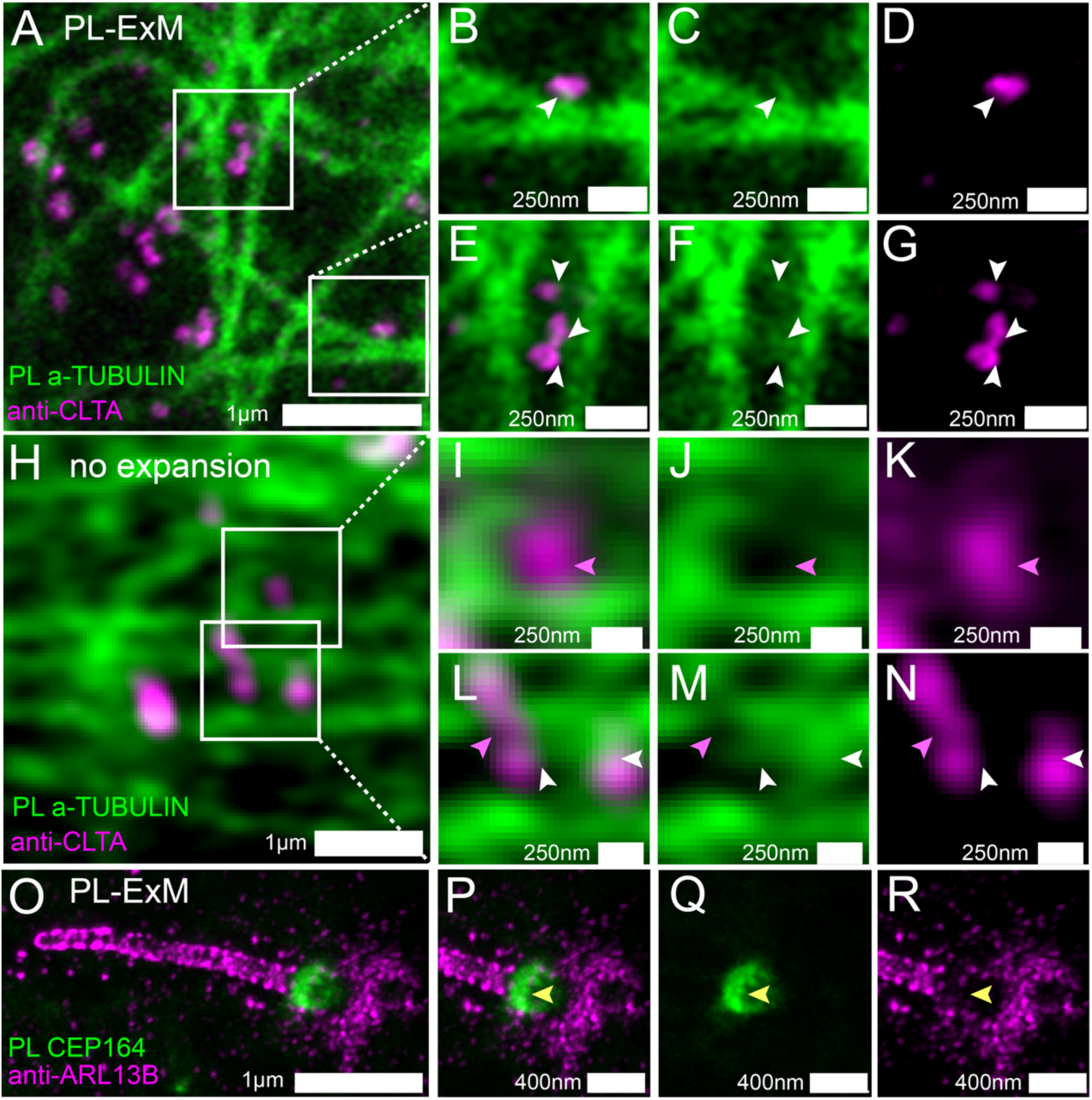
Two-color PL-ExM images dissect spatial relationships between interactive proteins. (A-G) PL-ExM images of proximity-labeled α-TUBULIN (green) and immunostained CLTA (magenta) in U2OS cells. (B-G) Magnified view of the boxed regions in (A). The white arrows indicate the co-localization of CCPs and bud-like structures stemming from microtubules. (H-N) Airyscan images of proximity-labeled α-TUBULIN (green) and immunostained CLTA (magenta) in U2OS cells without expansion. (I-N) Magnified view of the boxed regions in (H). The pink arrows point at CCPs that do not co-localize with microtubules. White arrows indicate possible colocalization of CCPs and microtubule structures. (O-R) PL-ExM of proximity-labeled CEP 164 (green) and immunostained ARL13B (magenta) in a primary cilium of a MEF cell. (P-R) Magnified view of the ciliary base in (O). The yellow arrows indicate anti-localization between ARL 13B and CEP 164. (A-N) are single-slice images. (O-R) are maximum intensity projections of z stacks. The length expansion factors are 4.1 (A-G) and 8.4 (O-R). All images are taken by an Airyscan microscope. All scale bars are in pre-expansion units.

We further applied PL-ExM on the primary cilium, a more challenging organelle with less abundant and tiny size (Figures 3O-R). The primary cilium is a sensory organelle that organizes signaling pathways, such as sonic hedgehog signaling, and their regulatory GTPases, such as ADP-ribosylation factor-like protein 13B (ARL13B). Mick et al. developed a groundbreaking method called cilia-APEX, which proximity-labeled ciliary interactome or MS analysis^45^. Using this method, they identified new components of cargos transporting GPCRs in cilia. Here, our aim is to use PL-ExM as a complementary method to cilia-APEX proteomics, providing spatial information. In this demonstration, we investigated a specific question: do the distal appendages (DAs) located at the base of the cilium mediate ARL13B entry or exit from the primary cilium? We simultaneously imaged proximity-labeled DA component CEP164 and immunostained ARL13B in MEF cells, using the two-color PL-ExM. With an 8.4-time expansion, we were able to resolve the donut-shaped DA disk and the distribution of AL13B through the cilia (Figure 3P). The images showed negligible overlapping between the interactome of CEP164 and ARL13B (Figures 3Q&R). The results indicated that the ARL13B either has no interaction or has very transient interaction with DAs.

### PL-ExM assesses the resolution and efficiency of proximity labeling

Despite PL’s capability of labeling interactomes, the labeling resolution and efficiency vary in each experiment. The parameters that cause the variability include the choice of enzyme, such as HRP and APEX2, the choice of labeling probes, such as different phenols, as well as the labeling duration^1, 46^. In this section, we will demonstrate how PL-ExM assesses PL under different enzymes (APEX2 vs HRP) and durations (30 seconds vs 20 minutes). We evaluated the quality of PL in each condition based on two important characteristics: labeling resolution and efficiency. The labeling resolution determines the spatial selectivity of the interactome and positive false rates, while the labeling efficiency indicates the coverage of the interactome. We used the average mitochondrial diameter (n≥90) measured from PL-ExM images as the readout of labeling resolution and total fluorescence intensity to compare the labeling efficiency between PL conditions. For fair comparison, all samples to be compared were labeled in the same batches (n>3) and imaged under the same microscope settings on the same days.

We compared two commonly used enzymes, APEX2 and HRP using PL-ExM. Mitochondrial outer membrane proteins were chosen as the bait proteins because their interactomes were extensively studied with PL-MS^10, 47^. The proteomic data can be used as references to validate our PL-ExM assessment. APEX2-catalyzed PL was performed on U2OS cells overexpressing *APEX2-OMM* (Figure 4A), where OMM is a peptide on the outer mitochondrial membrane. HRP-catalyzed PL was performed on U2OS cells which had TOMM20 immunostained with HRP-conjugated antibodies (Figure 4C). The same biotin-phenol and reaction duration were given in HRP and APEX2-cataluyzed PL. PL-ExM showed that the HRP-catalyzed PL achieved about four times higher labeling efficiency than the APEX2 condition (Figures 4B,D&E). In addition, The PL catalyzed by HRP also exhibited higher labeling resolution than APEX2, showing a smaller mitochondrial diameter of 0.56μm ± 0.030μm. (Figure 4G). On contrary, APEX2-catalyzed PL showed more diffusive signal around mitochondria (Figure 4A), resulting in a bigger mitochondrial diameter of 0.97μm ± 0.065μm (Figure 4F). The lower labeling efficiency and lower labeling resolution of APEX may be attributed to the limited permeability of biotin-phenol in live cells and the lower catalytic activity of APEX compared with HRP.

**Figure 4.**
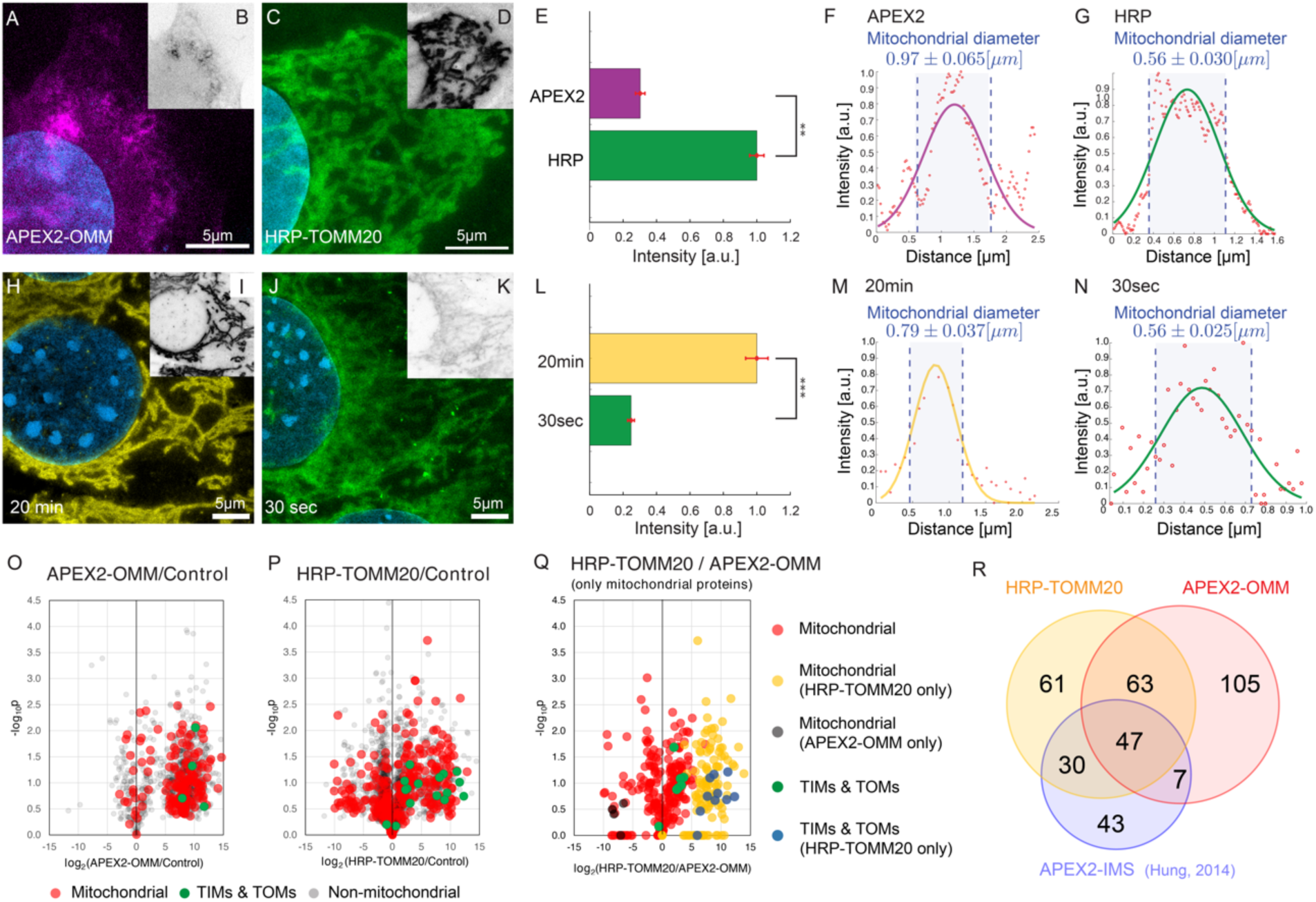
PL-ExM evaluates the labeling resolution and efficiency of APEX2- and HRP-catalyzed PL. In the comparison between APEX2 and HRP (A-N and O-R), APEX2-catalyzed PL was performed on U2OS cells overexpressing *APEX2-OMM*. HRP-catalyzed PL was performed on U2OS cells which had TOMM20 immunostained with HRP-conjugated antibodies. All images were taken on a confocal microscope with the same imaging condition. (A) Representative PL-ExM image of APEX2-catalyzed PL. (C) Representative PL-ExM image of HRP-catalyzed PL. (B,D) Grayscale images of A and C respectively. Brightness and contrast are set the same for these two images for the quantitative comparison. (A-D) are maximum intensity projections of 3D z-stacks for the same z depth. (E) The bar chart summarizes the fluorescence intensity of PL-ExM images of APEX2 and HRP samples. n ≥3 per condition. The reported p-value is smaller than 0.01. (F) A representative histogram showing the fluorescence intensity in a cross-section of a mitochondrion from a PL-ExM image of an APEX2 sample. The measured mitochondrial diameter is 0.97 ± 0.065μm. The mean and a standard error were obtained from 90 measurements across 3 independent samples. (G) A representative histogram showing the fluorescence intensity in a cross-section of a mitochondrion from a PL-ExM image of a HPR sample. The measured mitochondrial diameter is 0.56 ± 0.030μm. The mean and standard error were obtained from 90 measurements across 3 independent samples. In the comparison between 20-minute and 30-second reaction duration (H-N) HRP-catalyzed PL was performed on MEF cells that had TOMM20 immunostained with HRP-conjugated antibodies. (H) Representative PL-ExM image of HRP-catalyzed PL with 20-minute H_2_O_2_ treatment. (J) Representative PL-ExM image of HRP-catalyzed PL with 30-second H_2_O_2_ treatment. (I, K) Grayscale images of H, J respectively. Image brightness and contrast are set to be the same for the quantitative comparison. (A-K) Images are maximum intensity projections of 3D z stacks for the same z depth. (L) Labeling efficiency comparison between samples with 20-minute and 30-second H_2_O_2_ treatment. 20-minute samples show ∼4 times higher labeling efficiency than 30-second samples with p-value smaller than 0.001. The bar chart summarizes the fluorescence intensity of PL-ExM images from 20-minute and 30-second samples. n ≥3 per condition. (M) A representative histogram showing the fluorescence intensity in a cross-section of a mitochondrion from a PL-ExM image of a 20-minute sample. The measured mitochondrial diameter is 0.79 ± 0.037 μm. The mean and standard error were obtained from 90 measurements across 3 independent samples. (N) A representative histogram of a 30-second sample. The measured mitochondrial diameter is 0.56 ± 0.025 μm. The mean and standard error were obtained from 90 measurements across 3 independent samples. (O-P) Volcano plots depicting protein enrichment by APEX2-OMM (O) and HRP-TOMM20 (P). Fold-change is represented in log2 along x-axis, calculated as the relative normalized abundances of proteins in labeled/control. Subunits of the TIM/TOM complex are shown in green, while other mitochondrial proteins defined by MitoCarta are shown in red. Non-mitochondrial proteins are shown in gray. (Q) Mitochondrial protein enrichment by APEX2-OMM versus HRP-TOMM20. Log2 fold-change is represented along x-axis, calculated as the relative normalized abundances of proteins from HRP-TOMM20/APEX2-OMM. TIM/TOM complex subunits quantified by both APEX and HRP labeling shown in green, while those only quantified by HRP are shown in blue. The remaining mitochondrial proteins are shown in red, unless only quantified by APEX (black) or HRP labeling (blue). (R) Overlaps of enriched mitochondria proteins by APEX2-OMM, TOMM20-HRP, and APEX2-IMS^10^. The length expansion factors of PL-ExM images (A, C, H, J) are 4.1 ∼ 4.2. All scale bars are in pre-expansion units, and they are 5µm.

We also evaluated the PL quality with two labeling durations: 30 seconds and 20 minutes (Figures 4H-N). HRP was used to proximity label the TOMM20 in both conditions. The only difference is the duration of H2O2 treatment. We observed a nearly quadrupled labeling efficiency in the 20-minute condition, compared with the 30-second condition (Figure 4L). However, the diameter of the mitochondria measured from the two conditions did not differ that much. PL-ExM images of the 20-minute group showed a considerably larger mitochondrial diameter (0.79µm, Figure 4M), compared with 0.56µm of the 30-second group (Figure 4N). These results indicate the labeling efficiency of HRP-catalyzed PL significantly increases over time, while the labeling resolution drops only slightly. This finding underscores the importance of the PL treatment duration as a crucial variable that requires meticulous calibration based on the research objective.

To assess PL-ExM accuracy, we compared PL-MS and PL-ExM results from identically prepared samples as described above. The cells biotinylated by APEX2 and HRP were lysed, affinity purified, digested, and analyzed by MS. In comparison to non-labeled controls, label-free based quantitative MS analyses revealed that both APEX2 and HRP methods were able to enrich mitochondrial proteins (Figures 4O&P), which are comparable to a previous report using APEX2-IMS (Figure 4P) ^10^. Interestingly, HRP samples yielded stronger labeling of TIMs and TOMs proteins than the APEX2 samples, suggesting HRP-catalyzed PL was less diffusive and more effective in labeling proteins in closer proximity to the bait (TOMM20) (Figure 4Q). This observation is in good agreement with PL-ExM images. In summary, PL-ExM emerges as an invaluable tool in ascertaining the optimal experimental conditions for PL.

### PL-ExM is compatible with tissues

In previous sections, we have demonstrated the compatibility of different PL-ExM cell lines, such as U2OS and MEF used in Figures 1-4. Here, we move forward to apply PL-ExM to tissues. Since live cell PL is usually not applicable to tissues, we recommend the HRP-catalyzed PL approach of PL-ExM for interactome visualization for tissues. This way, HRP is tagged to the protein of interest in fixed tissue samples by antibodies. As a showcase, we applied HRP PL-ExM to mouse brains expressing neuron-specific marker Thy1 with YFP. We proximity-labeled Thy1 in the brain sections using the HRP approach (see Methods for more details). The x4 PL-ExM images displayed the distribution of the proximity-labeled interactome of protein Thy1 across the brain section (Figure 5A). Compared with cultured cells, the noise level of PL of tissues was higher. However, individual dendrites and axons of neurons can be clearly seen in the PL channel (green in Figure 5B). Furthermore, we co-immunostained an astrocyte marker Glial fibrillary acidic protein (GFAP) in the brain tissue. The two-color images showed the spatial entanglement between astrocytes and neurons, indicating their interactions (Figure 5B).

**Figure 5.**
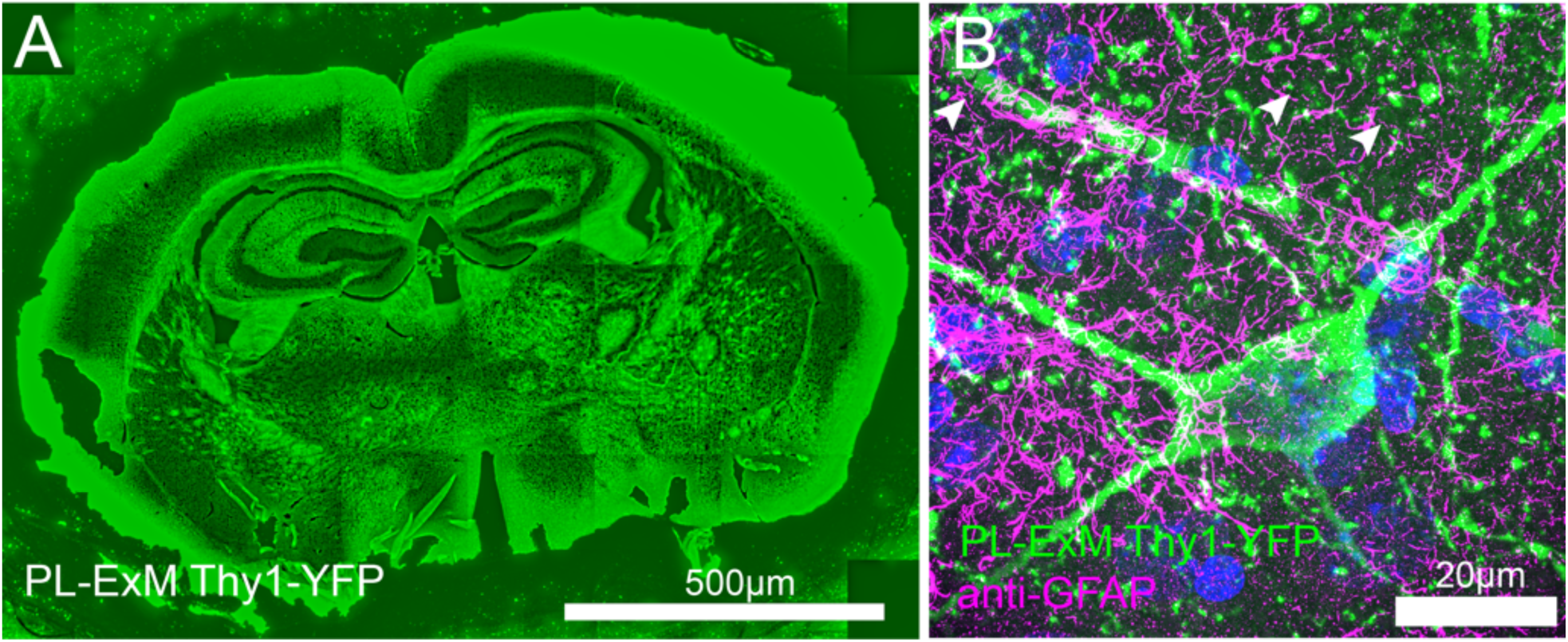
Two-color PL-ExM imaging reveals interactions in mouse brain tissues. Both images are Airyscan PL-ExM images of 20-µm sections of a mouse brain expressing Thy1-YFP with proximity-labeled Thy1-YFP (green) and immunostained GFAP (magenta). (A) Proximity-labeled Thy1-YFP channel of a whole mouse brain slice with. (B) A magnified view of (A) with both proximity labeled Thy1-YFP (green) and immunostained GFAP (magenta). Both mages are maximum intensity projections of z stack. The scale bars are 500 µm for (A) and 20µm for (B). The length expansion factor is 4.0. All scale bars are in pre-expansion units.

Deep imaging of tissue samples poses inherent challenges owing to the light scattering between layers of cell and extracellular matrix. The expansion procedure of PL-ExM transforms the intact tissue into a hydrogel that is optically transparent, sharing the same clearing principle with CLARITY ^48^. Therefore, PL-ExM offers tissue clearing for more clear and deeper visualization of the tissue structure, in addition to the super resolution.

## DISCUSSION

During the expansion procedure of PL-ExM, the homogenization step breaks down protein-protein interactions and the hydrogel expansion pulls interacted proteins away. There might be a question: will the breakdown of protein-protein interactions cause incomplete interactome detection in the images? The answer is no. It is because the interactome is defined by the PL, not the expansion. Proteins within the labeling radius are marked by biotin during the PL reaction when the cells are intact before the expansion procedure. Therefore, as long as the biotin signal can be detected after expansion, the breakdown of protein-protein interactions during the expansion procedure will not cause incomplete interactome detection. The highly efficient detection of biotin after expansion was proved by LR-ExM method that we recently developed ^34^.

The next question is about the fidelity of expansion. If the expansion is anisotropic, distortion of the interactome structure could happen during the expansion step, resulting in unreliable observation. Our team, along with other ExM developers, have rigorously ensured isotropic expansion, with optimization of fixation methods, protein anchoring efficiency, sample homogenization, and hydrogel recipes ^22, 42, 49, 50^. We have comprehensively discussed the solutions to make isotropic expansion of different biological samples in a recent review ^42^. This PL-ExM method is optimized for faithful expansion of proximity-labeled samples with different enzymes and reaction conditions. Either MA-NHS, glutaraldehyde, or glycidyl methacrylate worked well for the anchoring of biotinylated proteins. Like other ExM protocols, proteinase K digestion is a reliable sample homogenization method in PL-ExM. In quantitative comparison of different PL methods, it is important to apply the same anchoring and homogenization reagents and conditions to each sample.

In this work, we demonstrated 22 nm resolution by expanding cells 8.2 times using the TREx protocol ^38^ and imaging on an Airyscan microscope (Figures 2M-R). Higher resolution of PL-ExM can be achieved with up to 20 times expansion^35-40^ and a more advanced microscope, such as PALM, STORM, and STED. However, there is an upper limit to how high the resolution can be achieved using PL-ExM. Technically, the ultimate resolution is constrained by the pore size of the hydrogel before expansion. Because the pore size determines how fine the hydrogel can faithfully anchor the biomolecules in their initial positions. Any structural details smaller than the pore size are distorted.

During the method development, we found that the variabilities of PL labeling quality was often overlooked. The super-resolution of PL-ExM allowed us to directly observe the variation. The PL quality not only varied between methods, but also was influenced by the condition of the samples and human errors. The high concentration of radical quenchers in the cytosol and mitochondrial matrix^51^, along with macromolecular crowding^52^, could impact the spatial resolution and efficiency of PL^1^. It is important to note that biological systems are inherently variable and dynamic, influenced by genetics, environmental conditions, or the physiological state of the sample, which can introduce variability into the outcomes of PL. Therefore, we strongly recommend the developers of PL methods use super-resolution imaging, such as PL-ExM, to characterize the new methods. Similarly, we recommend PL-MS users to assess their sample preparation with PL-ExM. The spatial information provided by PL-ExM will aid in interpreting proteomic results and ruling out false positives.

## CONCLUSIONS and FUTURE DIRECTIONS

PL-ExM significantly advances interactome imaging by uncovering the intricate spatial organization of proteins within the interactome structure. By integrating the spatial biotinylation of interactive proteins throughout the PL with the enhanced imaging resolution offered by ExM, this method provides up to 12 nm resolution using conventional microscopes, including confocal and Airyscan. Our study showcased the potential of two-color PL-ExM by imaging the interactome in mitochondria, microtubules, clathrin-coated pits, and primary cilia. The results revealed detailed spatial organization of specific proteins within the context of the interactome architecture. The PL-ExM, which provides 3D structural information of the interactome, can be used as a complementary tool to the PL-MS interactome analysis. As we look to the future, the next frontier for PL-ExM would be to expand its multiplexity beyond the current two-color limitation. By incorporating highly multiplexed immunostaining techniques, like Immuno-SABER^53^, PL-ExM holds the promise of mapping every individual protein within the interactome. Ultimately, the true power of PL-ExM lies in its potential to unearth previously undiscovered 3D spatial relationships between interactive proteins, paving the way for a deeper understanding of intricate biological and pathological processes.

PL-ExM also stands out as a pivotal tool for gauging both the labeling resolution and efficiency of PL Methods. Our evaluation of APEX2- and HRP-catalyzed PL methods showed that PL-ExM has the resolving power to measure the labeling radius and has the sensitivity to compare the labeling efficiency across different PL methods. PL-ExM is compatible with a broad spectrum of PL methods that biotinylate proteins, including but not limited to APEX, HRP, BioID, TurboID, and µMap. The congruence between our imaging findings and the proteomic outcomes from PL-MS confirmed PL-ExM as a reliable quality control method for PL methodologies.

## METHODS

### Cell line generation

*APEX2-OMM* gene fragment (from a plasmid Addgene #238450) was cloned into a second generation 5’ self-inactivating lentiviral backbone (pHR) downstream of a SFFV promoter, using InFusion cloning (Takara Bio #638910). A pantropic VSV-G pseudotyped lentivirus was produced via transfection of Lenti-X 293T cells with the pHR transgene expression vector and viral packaging plasmids pCMVdR8.91 and pMD2.G using Fugene HD (Promega #E2312). At 48 hours, the viral supernatant was harvested, filtered through a 0.45 μm filter (Millipore #HAWP04700), and added onto the U2OS cells for transduction. *APEX2-OMM* cell lines are generated from Single-cell cloning of the transduced U2OS cells.

### Cell culture

MEF cells were cultured in DMEM, Glutamax (Thermofisher; 10566-016) supplemented with 15% Fetal Bovine Serum (FBS) and 1% antibiotics antimycotic solution (Sigma Aldrich; A5955) at 37°C in 5% CO2. U2OS (ATCC; HTB-96) and U2OS-*APEX2-OMM* cells were cultured in McCoy’s 5a (ATCC; 30–2007) supplemented with 10% FBS and 1% antibiotics antimycotic solution at 37°C in 5% CO2. For PL-ExM, cells were seeded at 10^4^ cells/cm^2^ in 16-well chambers (Grace Bio-Labs; 112358) and grown to 80% confluency. For MEF cells, we coat the chamber with gelatin solution (Sigma-Aldrich; G1393-100ML) for 1 hour at 37°C. In Figure 4O-R, MEF cells were seeded at a density of 10^4^cells/cm^2^ in 16-well chambers. After 16 hours of incubation, cells were starved for 24 hours in Opti-Mem reduced serum medium for ciliation.

### Animal Sacrifice and brain slice preparation

Thy1-YFP mice were euthanized via CO2 inhalation and transcardially perfused with ice-cold 1X PBS buffer. Brains were removed carefully and fixed in freshly made 4% paraformaldehyde solution for 24 hours at 4°C. Brains were then cryoprotected in 30% sucrose solution at 4°C before embedding in OCT and storage at -80°C. Frozen brains were sectioned at 20 μm on a Leica SM2000 R sliding microtome for subsequent immunohistochemical analyses. All animal protocols were approved by the Institutional Animal Care and Use Committee (IACUC) of the University of California, Irvine.

### HRP antibody catalyzed PL for cultured cells

Fixation, endogenous peroxidase blocking, permeabilization, and endogenous biotin blocking. In figure 2, MEF cells were fixed with 3% paraformaldehyde (PFA) and 0.1% Glutaraldehyde (GA) solution for 15 minutes at room temperature, followed by reduction using 0.1% sodium borohydride in PBS for 5 minutes. In Figure 4 A-N, cells were fixed with 3.2% PFA in PEM buffer (100 mM Pipes, 1 mM EGTA, and 1 mM MgCl2, pH 6.9) at room temperature for 10 minutes, followed by reduction using 0.1% sodium borohydride in PBS for 5 minutes. In figure 4O-R, cells were fixed with 4% PFA for 15 minutes at room temperature.

After fixation, cells were washed with PBS for 3 times, with 5 minute interval between washes. Then, cells were incubated with 3% hydrogen peroxide (H2O2, Sigma Aldrich; H1009) for 5 minutes at room temperature to block the endogenous peroxidase before introducing any HRP in the system. Reaction was quenched by adding 2mM of L-Ascorbic acid sodium (Alfa Aesar; A17759) for 5 minutes followed by three PBS wash. The fixed cells were incubated in a permeabilization/blocking buffer (3% BSA, and 0.1% Triton X-100 in PBS) for 30 minutes at room temperature prior to immunostaining steps.

Primary antibodies at a concentration of 2 µg/ml were added to the fixed cells in the blocking buffer (3% BSA in PBS) for 16 hours at 4°C. The primary antibodies used for this paper are Rabbit x TOMM20 (1:250 dilution, santa cruz; sc-11415), Rat x α-TUBULIN, tyrosinated, clone YL1/2 (Millipore Sigma; MAB1864-I), Rabbit x anti-clathrin heavy-chain (1:100 dilution, Abcam; ab21679), Rabbit x ARL 13B (1:100 dilution, Proteintech; 17711-1-AP), Mouse x CEP164 (1:100 dilution, Santa Cruz; sc-515403), Chicken x GFAP (1:1000 dilution, AbCam; ab4674), Rabbit x GFP (D5.1,1:200, Cell Signaling; 2956). After primary antibody incubation, the cells were washed with a blocking buffer for three times followed by 5 minutes of incubation between washes. After washing, cells were incubated with 3 µg/mL AffiniPure Goat x Rabbit (1:100, Jackson ImmunoResearch; 111-005-144), Goat x Mouse (1:100, Jackson ImmunoResearch; 115-005-146), or Goat x Rat (1:100, Jackson ImmunoResearch; 112-005-167) secondary antibodies in blocking buffer for 1hour at room temperature, then the cells were washed with a blocking buffer for three times followed by 5 minutes of incubation between washes. After secondary antibody staining and washing, cells were incubated with ImmPRESS HRP Horse x Goat (no dilution, Vector Laboratories; MP-7405) for 1 hour followed by three washing with PBS.

Cells were incubated with 0.5mM biotin phenol solution (Biotin tyramide, Sigma Aldrich; SML-2135) for 15 minutes at room temperature. A fresh 2mM H2O2 solution (in PBS) was prepared right before the reaction, and the same volume of H2O2 solution was added to the cells in the biotin phenol solution for 30 seconds if specified otherwise. After treatment, the reaction was quenched with 2mM of L-Ascorbic acid sodium solution for 5 minutes at room temperature.

### APEX2-catalyzed PL for cultured cells

Permeability of biotin phenol has significant implications on the efficacy of proximity labeling, emphasizing the need for careful calibration when proximity labeling is done when cells are live. We tested 1mM biotin phenol incubation for 2 hours at 37°C gives the best labeling results. A fresh 2mM H2O2 solution (in PBS) was prepared right before the reaction, and the same volume of H2O2 solution was added to the cells in the biotin phenol solution for 1 minute. After treatment, the reaction was quenched with 2mM of L-Ascorbic acid sodium solution for 5 minutes, followed by three PBS washes. After proximity labeling, U2OS-*APEX2-OMM* cells were fixed with 4% PFA for 15 minutes at room temperature and washed with PBS for 3 times.

### HRP antibody catalyzed PL for mouse brain tissues

We first dried a tissue slide for 30 minutes and rehydrated it for 10 minutes by immersing the sample in PBS. After additionally washing the sample with PBS for 2 times, we incubated a tissue sample with 3% hydrogen peroxide for 5 minutes. The reaction was quenched by adding 2mM of L-Ascorbic acid sodium and incubating for 5 minutes followed by PBS wash for three times. Then the tissue sample was incubated in a permeabilization/blocking buffer (3% BSA, and 0.1% Triton X-100 in PBS) for an hour. We performed overnight primary antibody staining at 4°C using Rabbit x GFP (D5.1,1:200, Cell Signaling; 2956), followed by 2.5 hour of Goat x Rabbit secondary antibody staining (1:100, Jackson ImmunoResearch; 111-005-144), and 2.5 hour of tertiary staining using ImmPRESS HRP Horse x Goat (no dilution, Vector Laboratories; MP-7405). After series of antibody staining, we incubated tissue sample in 0.5mM biotin phenol solution (Biotin tyramide, Sigma Aldrich; SML-2135) for 15 minutes. A fresh 2mM H2O2 solution (in PBS) was prepared right before the reaction, and the same volume of H2O2 solution was added to tissue sample in the biotin phenol solution for 30 seconds for proximity labeling. After treatment, the reaction was quenched with 2mM of L-Ascorbic acid sodium solution for 5 minutes. After proximity labeling step, we performed additional immunostaining on GFAP for 2.5 hours using primary antibody Chicken x GFAP (1:1000 dilution, AbCam; ab4674). Then we performed secondary antibody staining for 2.5 hours using Donkey x Chicken Dig-MA-NHS (prepared in our lab). After immunostaining, we performed anchoring for 10 minutes using 0.25% glutaraldehyde solution. Tissue sample was gelated, stained and expanded in a similar way to the Label-Retention expansion microscopy^34, 41^. All reactions are done at room temperature, and after each step sample was washed for 3 times in PBS (unless it is specified otherwise).

### Protein anchoring, gelation, denaturation, post-digestion fluorescent staining, and expansion steps of the x4 PL-ExM

Protein anchoring: After PL and immunostaining of the samples, one of the three anchoring reagents has been used: 0.25% Glutaraldehyde (GA; Electron Microscopy Sciences; 16120) solution prepared in PBS for 10-minute room temperature incubation, 25mM Methacrylic acid N-hydroxysuccinimide ester (MA-NHS; Simga-Aldrich; 730300) solution prepared in PBS for 1-hour room temperature incubation or 0.04% glycidyl methacrylate solution prepared in 100mM sodium bicarbonate, pH 8.5 (GMA; Sigma-Aldrich; 151238) for 4-hour room temperature incubation. The three anchoring reagents yielded similar anchoring efficiency.

Gelation, denaturation, fluorescent staining, and expansion have been performed in a similar way to the Label-Retention expansion microscopy (LR-ExM) ^34, 41^. Here we describe the procedure briefly.

Gelation: The samples were first incubated with monomer solution (8.6 g sodium acrylate, 2.5 g acrylamide, 0.15 g N,N’-methylenebisacrylamide (bis), 11.7 g sodium chloride in 100 ml PBS buffer) on ice for 5 min. Gelation solution (mixture of monomer solution, 10% (w/v) N,N,N′,N′ Tetramethylethylenediamine (TEMED) stock solution, 10% (w/v) ammonium persulfate (APS) stock solution and water at 47:1:1:1 volume ratio) was then quickly added to the samples and incubated on ice for another 5 min. The samples with gelation solution were later transferred to a 37 °C humidity chamber for gelation for 2 hours.

Denaturation: After 1 h gelation, the gelated samples were immersed in proteinase K buffer (8 units/mL proteinase K in digestion buffer made of 50 mM Tris pH 8.0, 1 mM EDTA, 0.5% Triton X-100, 1M NaCl), and then washed with excess of DNase/RNase-free water. For cultured cells, the proteinase K incubation duration was 16 hours at room temperature. For tissues, the duration was 1.5 hours at 78°C.

Post-digestion fluorescent staining: The gelated samples were incubated in a mixture of 3 uM fluorescently labeled streptavidin (e.g. streptavidin-Alexa Fluor 488) and fluorescently labeled anti-DIG antibodies (e.g. anti-DIG-DyLight 594) buffer for 24 hours at room temperature. The staining buffer comprises 10 mM HEPES and 150 mM NaCl in water at pH 7.5.

Expansion: The gelated samples were expanded in DNase/RNase-free water for more than 4 hours at room temperature. Fully expanded gelated samples were trimmed and transferred to a poly-lysine-coated glass bottom multiwell plate or dish for imaging.

### Protein anchoring, gelation, denaturation, post-digestion fluorescent staining, and expansion steps of the x8 PL-ExM

The anchoring, digestion, and post-digestion fluorescent staining steps of the x8 PL-ExM were identical to those of the x4 PL-ExM. The gel monomer recipe and expansion steps of the 8x PL-ExM were modified based on the TREx protocol^38^. Briefly, the samples were first incubated with monomer solution for x8 expansion (1.1 M sodium acrylate, 2.0 M acrylamide, 50 ppm bis in PBS) on ice for 5 min. Gelation solution (mixture of monomer solution,1.5 ppt APS, and 1.5 ppt TEMED) was then quickly added to the samples and incubated on ice for another 5 min. The samples with gelation solution were later transferred to a 37 °C humidity chamber for gelation for 2 hours. The expansion step was similar to that of the x4 PL-ExM except for the overnight expansion duration at room temperature.

### Image acquisition and analysis

Airyscan imaging for PL-ExM data was performed on Zeiss LSM 980 and Zeiss LSM 900 with a 63x water immersion objective (Zeiss Plan Apo 63x NA 1.15). Non-expanded samples were imaged with Airyscan mode using Zeiss LSM 980 with a 63x water immersion objective (Zeiss Plan Apo 63x NA 1.15). Confocal imaging was performed on either Zeiss LSM 980 using 63x water immersion objective (Zeiss Plan Apo 63x NA 1.15) or a spinning-disk confocal microscope (Nikon CSU-W1 Sora) with a 40× water-immersion objective (Nikon CFI Apo 40× WI NA 1.15). The fluorescence intensity of Airyscan and confocal images was analyzed using the open-source software Fiji (ImageJ). No deconvolution was applied to any images in this work.

### Image intensity quantitative analysis and statistics

Images were first denoised where we define a noise such as

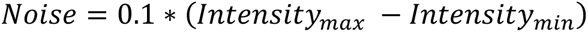

We use Matlab improfile function to select the cross-sectional area of proximity labeled diameter and fit the Gaussian function and measure the full width half maximum (FWHM) from it. We used single-slice images to measure the FWHM. Customized Matlab codes were used, and the codes are available upon request. The mean and a standard error were obtained from >=90 measurements across 3 independent samples. For Figure 4, student t-test was performed to calculate p-value and determine statistical significance.

### Protein purification and digestion for MS

The cell pellets were lysed in lysis buffer [50 mM Tris-HCl, 500 mM NaCl, 0.2% SDS, 1% Triton, 1 mM Tris(2-carboxyethyl) phosphine hydrochloride (TCEP), 10 mM sodium azide, 10 mM sodium ascorbate, 5 mM TROLOX, protease inhibitor cocktail (pH 7.5)] with sonication on ice. The lysates were centrifuged at 13, 000 rpm for 15 minutes to remove cell debris, and the supernatant was incubated with streptavidin Mag Sepharose resin (Cytiva) for overnight at 4°C with rotation. The streptavidin beads were then washed twice with four buffers containing: A) 2% SDS at room temperature; B) 50 mM Tris-HCl, 500 mM NaCl, 2% Triton-X; C) 50 mM Tris-HCl, 250 mM NaCl, 0.5% SDS, 0.5% Triton-X and D) 2 M Urea, 50 mM Tris-HCl at 4 °C. The bound proteins were then reduced, alkylated, and digested on-bead by LysC in 8M urea/25mM NH4HCO3 for 4 hours, followed by trypsin in 1.5 M urea/25 NH4HCO3 overnight at 37°C. The peptide digests were extracted and desalted with C18 tip (Agilent) prior to liquid chromatography tandem mass spectrometry (LC MS/MS)^54^.

### Mass spectrometry analysis

The peptide digests were subjected to LC MS/MS analysis using an UltiMate 3000 RSLC system (Thermo Fisher Scientific) coupled in-line to an Orbitrap Fusion Lumos mass spectrometer (Thermo Fisher Scientific). Reverse-phase separation was performed on a 50 cm x 75 μm I.D. Acclaim® PepMap RSLC column. Peptides were eluted using a gradient of 4% to 22% B over 87 minutes at a flow rate of 300 nL/min (solvent A: 100% H2O, 0.1% formic acid; solvent B: 100% acetonitrile, 0.1% formic acid). Each cycle consisted of one full Fourier transform scan mass spectrum (375–1500 m/z, resolution of 120,000 at m/z 400) followed by data-dependent MS/MS scans acquired in the Orbitrap with HCD NCE 30% at top speed for 3 seconds. Target ions already selected for MS/MS were dynamically excluded for 30s. Protein identification and label-free quantitation was carried out using MaxQuant as described ^55^. Raw spectrometric files were searched using MaxQuant (v. 2.0.3.0) against a FASTA of the complete human proteome obtained from SwissProt (version from April 2023). The first search peptide tolerance was set to 15 ppm, with main search peptide tolerance set to 4.5 ppm. Trypsin was set as the digestive enzyme with max 2 missed cleavages. Methionine oxidation and protein N-terminal acetylation were set as variable modifications, while cysteine carbamidomethylation was set as a fixed modification. Peptide spectra match and protein FDRs were both set as 0.01. For quantitation, intensities were determined as the full peak volume over the retention time profile. “Unique plus razor peptides” was selected as the degree of uniqueness required for peptides to be included in quantification. The resulting iBAQ values for each identified protein by MaxQuant were used for comparing protein relative abundances. For figure 3O-R, we performed two mass spectrometry experiments to make a quantitative comparison between PL performed on U2OS cells overexpressing APEX2-OMM vs PL performed on U2OS which has TOMM20 immunostained with HRP-conjugated antibodies. For each condition, we also included negative controls. First, we cultured both U2OS-*APEX2-OMM* (experimental, and negative control) and WT U2OS cells (experimental, and negative control) in multiple 150 mm dishes, trypsinized cells, and collected them into 1.5 mL Eppendorf tube after centrifugation at 1800 rpm for 3 minutes. Final counts used for each condition was about 2*10^8^ cells per condition. In figure 3O,Q,R, U2OS-*APEX2-OMM* cells were used. We treated both experimental and control conditions using 500µL of 1mM Bitoin Phenol solution (BP, in PBS) at 37°C for 2 hours. Without removing BP solution, the experimental condition was treated with the same volume of 2mM freshly prepared H2O2 solution for 1 minute, followed by the addition of 750µL of 15mM sodium ascorbate solution for reaction quenching. The sample was thoroughly washed using PBS for 2 times with each 3 minute interval. After the proximity labeling step, each sample was fixed with 1% paraformaldehyde (PFA) solution; the control condition was immediately fixed with freshly prepared 1% paraformaldehyde (PFA) after BP incubation (but no H2O2 treatment). After every step, we thoroughly homogenize the sample, and centrifuge the sample at 500G for 3 minutes to pallet the sample before next treatment. In figure 3P-R, WT U2OS cells were used. Cells were first fixed with 0.1 % glutaraldehyde (GA) for 15 minutes at room temperature, and then washed with PBS 3minutes for 3 times. We incubated cells with blocking buffer (3% BSA in PBS) for 30 minutes and performed primary antibody staining using Rabbit x TOMM20 (1:250 dilution, santa cruz; sc-11415) overnight at 4°C. After washing samples 3 times using blocking buffer (5 minute each), we stained samples with 3µg/mL AffiniPure Goat x Rabbit (1:100, Jackson ImmunoResearch; 111-005-144) in blocking buffer for 1hour at room temperature, then washed with blocking buffer three times (5 minute each). We then stained samples with Goat-HRP (no dilution, Vector Laboratories; MP-7405) for 1hour at room temperature, washed with blocking buffer 3 times for 5 minutes each. Next, we incubated cells in 500µL of 0.5mM BP solution at RT for 15 minutes. We stopped any further treatment to negative control at this step; meanwhile, the experimental condition was treated with 500µL of 2mM H2O2 solution for 30 seconds at room temperature, followed by the addition of 750µL sodium ascorbate solution. After 5 minute of incubation, samples were thoroughly washed with PBS 3 times.

### Image resolution measurement

0.1µm size fluorescent beads (TetraSpeck Microspheres, Invitrogen; T7279) were used to measure the resolution of the Airyscan LSM980 resolution with 63x water immersion objective (NA1.15). 30 different beads were sampled to obtain the average full width half maximum (FWHM) with standard error. Effective resolution of PL-ExM was measured by calculating FWHM divided by the physical expansion factor of the hydrogel.

## AUTHOR CONTRIBUTIONS

S.P. and X.S. conceived and led the research. S.P. performed PL-ExM, imaging, and prepared samples for MS analysis. X.W. performed MS experiments and analysis. X.L. and X.S. initialized the concept and performed preliminary experiments. X. H. made plasmids, and generated cell lines with X. S. K.F. performed cell experiments under S.P. supervision. A.A.T synthesized early versions of LR-ExM probes. Z.D. synthesized LR-ExM probes. L.S. assisted Airyscan imaging. L.H. led and supervised MS work. X.W performed MS experiments. X.W., C.Y and L.H analyzed MS data. K.F assisted sample preparation. S.P, X.S, and L,H. drafted and edited the manuscript.

### Notes

The authors declare no competing financial interest.

## ACKNOWLEDGMENT

We acknowledge Professor Vivek Swarup and Dr. Sudeshna Das for providing mouse brain tissues to this project. S.P. is supported by an NSF-Simons grant, DMS1763272 (594598). S.P and X.S. are supported by the K99/R00 NIH Pathway to Independence Award (K99/R00GM126136), the NIH Director’s New Innovator Award (DP2GM150017), and the Chan Zuckerberg Initiative (CZI) Visual Proteomics Imaging Award. X.W., C.Y., and L.H are supported by the NIH Maximizing Investigators’ Research Award (R35GM145249). Special thanks to the CZI Advancing Imaging through Collaborative Projects award for supporting our dissemination of PL-ExM and LR-ExM.

